# Precise Nanopore Signal Modeling Improves Unsupervised Single-Molecule Methylation Detection

**DOI:** 10.1101/2023.07.13.548926

**Authors:** Vladimír Boža, Eduard Batmendijn, Peter Perešíni, Viktória Hodorová, Hana Lichancová, Rastislav Rabatin, Broňa Brejová, Jozef Nosek, Tomáš Vinař

## Abstract

Base calling in nanopore sequencing is a difficult and computationally intensive problem, typically resulting in high error rates. In many applications of nanopore sequencing, analysis of raw signal is a viable alternative. Dynamic time warping (DTW) is an important building block for raw signal analysis. In this paper, we propose several improvements to DTW class of algorithms to better account for specifics of nanopore signal modeling. We have implemented these improvements in a new signal-to-reference alignment tool Nadavca. We demonstrate that Nadavca alignments improve unsupervised methylation detection over Tombo. We also demonstrate that by providing additional information about the discriminative power of positions in the signal, an otherwise unsupervised method can approach the accuracy of supervised models.

**Availability and implementation:** Nadavca is available under MIT license at https://github.com/fmfi-compbio/nadavca. Nanopore sequencing data sets are available from ENA bioproject PRJEB64246. *Jaminaea angkorensis* reference genome assembly is available from Zenodo https://doi.org/10.5281/zenodo.8145315.

## 1. Background

DNA sequencing devices developed by Oxford Nanopore Technologies, including the pocket-sized MinION, measure changes in electrical current as a DNA strand passes through a nanopore. The result is a sequence of signal measurements (also called *a squiggle*). In this paper, we provide an improved algorithm to align parts of the squiggle to a known reference DNA sequence and show that improved alignment accuracy benefits downstream analysis in de novo methylation identification.

Typically, the first step in the analysis of nanopore sequencing data is base calling, which translates a squiggle into a DNA sequence. The most successful base callers are based on complex machine learning models, such as recurrent neural networks [1, 2]. A typical median read accuracy achieved by a base caller is approx. 97% [3], but when base called reads are piled up at a deep coverage, the consensus accuracy can be as high as 99.94% [4].

An alternative class of approaches avoids base calling and works directly with the raw signal sequence. The main motivation is to use the additional information present in the raw signal, which is potentially lost in base calling. Applications include mainly better modification calling and methylation detection. Base calling is also computationally intensive, and thus in some situations, working with the raw signal can be faster.

### Basic properties of nanopore signal

The measured electrical current depends mostly on the context of *k* consecutive nucleotides passing through the nanopore. A signal model predicts for each context of size *k* the expected signal level. For the case of nanopores in R9.4.1 flow-cells, which were used in this paper, a typical model uses the context of size *k* = 6. The DNA moves through the pore at the speed of roughly 450 bases per second and the signal is read at a frequency of 4000 samples per second. This means that on average we have approx. 9 samples per one DNA context, but as we show in the results, the actual number of samples per context can vary significantly.

### Signal-to-reference alignment and dynamic time warping

One of the basic building blocks of nanopore signal analysis is the alignment of squiggles to a known reference sequence. The squiggle can be represented as a sequence of numbers *s*_1_…*s*_*n*_. For each nucleotide *r*_*i*_ of the reference sequence *r*_1_*r*_2_*… r_m_*, we need to assign the corresponding interval 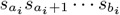 of the squiggle which we call an *event*. Such squiggle-to-reference alignment facilitates further analysis, allowing to find significant differences between the squiggle and the reference signal computed based on the signal model applied to the reference sequence. Identified differences typically correspond to sequence variants or modifications [5]. Alignments also enable pooling together information from multiple reads aligned to a single reference location and to visualize squiggles in the genomic context [6].

The squiggle alignment is typically performed by a class of algorithms called *dynamic time warping* (DTW), originating in the field of speech processing [7]. The alignment of a signal level *s* to a position *j* in the reference *r* is evaluated by a scoring function *d*(*s, r, j*), which typically takes into account the context around position *j*; for example, the scoring function *d*(*s, r, j*) = (*s−* expectedsignal(*r*_*j−*(*k−*1)/2_ *…r*_*j*+(*k−*1)/2_))^2^ uses a signal model based on the context of *k* nucleotides (assuming *k* is odd). Using a scoring function of this type, we are looking for the alignment minimizing the score 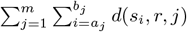, which can be computed using a dynamic programming algorithm similar to sequence alignment.

In our experience, the DTW-like algorithms applied in the context of MinION squiggle alignment show a variety of artefacts. The scoring function allows squiggle to be compressed or stretched arbitrarily in time, without any penalty. If several consecutive sequence contexts along the reference genome exhibit similar expected signal values in the signal model (which is not uncommon), there is a large uncertainty in how the corresponding observed signal should be split between the events. Depending on the exact DTW implementation, this may lead to artificially short or artificially long events. Small amounts of noise in the signal can easily lead to events split incorrectly between multiple reference contexts. In fact, the real squiggles often contain transitional readouts between signal levels corresponding to consecutive signal contexts, sometimes even alternating between two consecutive contexts for some time. If the squiggle does not correspond exactly to the reference sequence (e.g. in case of single nucleotide variants or base modifications), DTW tends to “hide” the differences between the reference and the squiggle within the neighboring events. This subsequently affects the ability to detect such differences.

In this paper, we propose several improvements to the DTW framework to overcome these obstacles. Instead of simple maximization of the scoring function, we propose to employ alternative methods similar to posterior decoding in probabilistic models [8, 9], to account for uncertainty in the event boundaries. We enhance the signal model underlying the scoring function to take into account a wider context of the sequence and also specifically model transitions between the contexts. We also employ a two-iteration approach to signal normalization to avoid artefacts resulting from incorrect shifting and scaling of the squiggle. We have implemented these improvements in an alignment tool called Nadavca, and we have evaluated their effect on the accuracy of detection of base modifications in nanopore reads. To make the running time practical, we use methods similar to banded and anchored sequence alignment.

### Supervised and unsupervised modification detection

It is known that modified nucleotides affect the signal levels when passing through nanopore [10]. For known modification patterns, such as CpG methylation, it is possible to prepare artificially methylated sequences and either estimate a signal model for contexts containing methylated nucleotides [11, 12, 13] or to train a machine learning algorithm, such as a neural network, to recognize signals from sequences that contain methylated nucleotides [14, 15, 16]. We call this approach *supervised methylation detection*, since substantial amounts of labeled training data are required to build such models.

In contrast, *unsupervised modification detection* can be used when labeled training data is not available, or when we are targeting previously uncharacterized DNA modifications. One option is to compare signal values in reads in the studied sample with signals in a control sample devoid of modifications and look for statistically significant differences [6, 17]. However, it is also possible to find putative modifications simply by comparing the actual signal with the expected signal predicted by a signal model trained on unmodified sequences (this approach is implemented for example in Tombo [18]). The strength of this method is that it requires only unmodified training data to obtain the signal model and then can be applied directly to biological samples from various organisms without the need for modified training data or matched unmodified dataset for each analysis. The crucial step in this approach is accurate alignment of signal to the reference to allow meaningful comparison with the model signal. Using Nadavca signal-to-reference alignments, we implemented a new tool nanometh for unsupervised modification detection. We demonstrate that Nadavca alignments increase the accuracy of unsupervised modification detection, and we also evaluate contribution of individual features implemented in Nadavca.

## 2. Methods

### 2.1. Realignment of Signal to the Reference with Posterior Decoding

The core part of Nadavca aligns a portion of nanopore signal (values *s*_1_,*…, s*_*n*_) to the corresponding part of the reference genome (bases *r*_1_, *…, rm*). The objective is to improve the accuracy of an approximate alignment, resulting from aligning base-called reads to the reference. The output alignment assigns an interval of the signal sequence 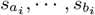 (also called event) to each base *ri* of the reference. For the basic version of the algorithm, we require that the intervals for adjacent bases are adjacent in the signal sequence (i.e. *a*_*i*_+_1_ = *b*_*i*_ + 1).

The standard dynamic time warping (DTW) algorithm is based on dynamic programming. We define subproblem *A*_*i,j*_, which represents the score of the best alignment of the squiggle segment *s*_1_*… s_i_* to the reference segment *r*_1_*… r_j_*, assuming that the signal *s*_*i*_ is aligned to the reference base *r*_*j*_. *A*_*i,j*_ can be computed gradually using a simple recurrence *A*_*i,j*_ = *d*(*s*_*i*_, *r, j*) + min(*A*_*i−*1,*j*_^*′*^, *A*_*i−*1,*j−*1_), where *d*(*s*_*i*_, *r, j*) is the score of aligning signal value *s*_*i*_ to the reference sequence *r* at position *j*. After finishing the computation, the final alignment (intervals [*a*_*i,bi*_]) can be easily computed using standard methods for tracing back the dynamic programming solution. The alignment can thus be visualized as a path in this dynamic programming matrix.

Our alignment algorithm is inspired by the posterior decoding approach to sequence alignment [8, 9]. While the standard DTW seeks the alignment with the highest overall score, our goal is to consider also a contribution of sub-optimal alignments. Many of these alignments can have scores very close to the optimum, representing uncertainty in the true alignment. Posterior decoding algorithms consider this uncertainty at each position of the alignment.

To apply this principle to DTW, we first modify the scoring scheme and instead of the distance score *d*(*s*_*i*_, *r, j*), we use probabilistic score *p*(*s*_*i*_, *r, j*), which gives a probability of signal value *s*_*i*_ being produced from the reference *r* at position *j*. Details of this probabilistic scoring scheme are described below. Now the score of the alignment becomes the product of probabilities along the alignment path 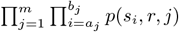. Higher scores indicate higher alignment quality.

The key concept in posterior decoding is the posterior probability *P*_*i,j*_ which for each *i* and *j* sums the scores of all alignments that align signal value *s*_*i*_ to the reference base *rj*. All values of *P*_*i,j*_ can be computed by Forward-Backward algorithm similarly as in the context of pair hidden Markov models [8, section 4.4]. Finally, we compute the resulting alignment as a path through the matrix *P* maximizing the product of individual posterior scores *P*_*i,j*_ (as in [9]), using a dynamic programming recurrence similar to the standard DTW.

In the real data applications, we impose additional restrictions. The minimum length of each event *b*_*i*_− *a*_*i*_ + 1 is constrained by the parameter *min_event_length*. The very start and the very end of the signal may remain unaligned. The length of unaligned portions is at most 2*bandwidth* + 1, where *bandwidth* is a parameter described below. Corresponding modifications of the algorithms are straightforward.

### 2.2. Enhanced Signal Model

Accurate models of signal levels expected in different sequence contexts are important for precise sequence alignment. Tombo [18] by default uses the 6-mer model supplied by Oxford Nanopore, which predicts the expected signal based on the sequence context of length 6. Instead, we use a (restricted) 10-mer model which we compute in the following way. First, we align the training dataset (described below) to the reference genome using Nadavca with the default 6-mer model. Computing the mean signal for each 10-mer would lead to a huge model, requiring a gigantic amount of data in order to cover each 10-mer with enough samples. Therefore, we restrict our 10-mer model by expressing it as the sum of three separate sub-models: a base 6-mer model *MC* for the central part of the context and two 4-mer models *M*1 and *M*_2_ that fine-tune the result based on more distant bases. For a context *r* = *r*_1_*r*_2_ … *r*_10_, the expected level μ(*r*) of the signal is computed as follows: μ(*r*) = *M*_1_(*r*_1_ … *r*_4_)+*MC* (*r*_3_ … *r*_8_)+*M*_2_(*r*_7_ … *r*_10_) (see Figure 1 for illustration). Models *M*_1_, *M*_2_, and *MC* are estimated simultaneously for all 4-mers and 6-mers respectively, by fitting the observed signal levels using the least-squares method. The probabilistic score *p*(*s*_*i*_, *r, j*) is now computed based on *s*_*i*_ ∼*N*(μ(*r*_*j−4*_ … *r*_*j+5*_), σ) for constant σ = 0.3.

**Figure 1:**
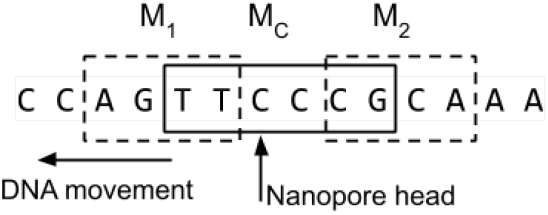
Extended signal model

### 2.3. Transitional States

The simplest models [19] of the nanopore signal assume that the signal can be partitioned into events, each event corresponding to one position in the reference and thus having a roughly uniform signal level influenced by the local sequence context. However, transitions between events are not instantaneous, and as a result, the measured signal may include intermediate values between the expected levels typical for two adjacent events or it may even jump back and forth between these levels (see Figure 2).

**Figure 2:**
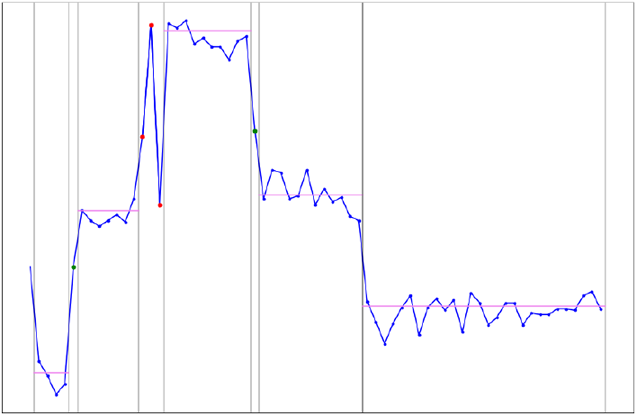
Example of transitions between states. Green dots show observed intermediate values between signal levels. Red dots represent signal values that jump back and forth between adjacent states.

In Nadavca, we handle these phenomena by inserting a special “transitional state” between each pair of adjacent reference bases *r*_*i*_ and *r*_*i+1*_. The primary score for aligning signal *s*_*j*_ to this transitional state is computed from a special distribution as opposed to the *k*-mer distribution used normally. While this distribution can depend on the sequence context, for simplicity we currently use a constant score 0.01. Thus, any signal aligned to these positions is penalized, regardless of the signal level and the sequence context. This simple heuristic works well in practice.

The posterior decoding algorithm aligns transitional states to intervals of signal values, but these intervals are not constrained to have a minimum required length; they can even be empty. The intervals aligned to transitional states are omitted from Nadavca’s output, and thus the output events corresponding to the reference bases may have gaps between them.

### 2.4. Fast Implementation With Banded Alignment

To avoid slow computation of the full dynamic programming matrix, we use the banded alignment heuristic around an approximate guide alignment obtained by aligning the base called sequence to the reference.

In particular, we first base call each read, and align the base called sequence to the target genomic sequence using BWA MEM [20]. The BWA MEM alignment will determine the window of the target genome to which we need to align the signal. This window is padded with several additional bases from base call to introduce context for the signal model; please note that these bases may be different from the target due to the adapters or barcodes present at the beginning or at the end of the read (see Figure 3). The resulting sequence will serve as the reference for the signal alignment.

**Figure 3:**
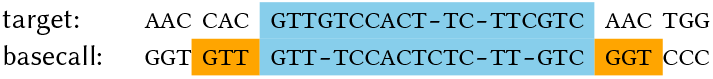
Alignment of the basecall to the target determines the reference window that needs to be aligned to the signal. Blue color highlights the aligned window, orange represents the context needed for the signal model.

In the next step, we consider all matched bases from the BWA MEM alignment. For each base called base *b*_*j*_ matched to a reference base *r*_*i*_, we locate the interval *s*_*c*…*d*_ in the signal that corresponds to *b*_*j*_ according to the Events table provided by Guppy. Interval *s*_*c*…*d*_ will be anchored to the reference base *r*_*i*_ and we call the set of all anchors the *guide alignment*.

We use the guide alignment to determine the portion of the signal that needs to be aligned to the reference. It will extend from the start of the interval in the first anchor to the the end of the interval in tha last anchor, extended further on both ends by the *bandwidth* parameter.

The guide alignment will also define an area of interest for dynamic programming. Recall that our posterior alignment uses three passes of dynamic programming (forward, backward, and posterior), and all three of them will consider only such alignment paths that are contained within the area of interest. All values outside of the area of interest are considered as zeroes. For each reference base *r*_*i*_, the area of interest contains the interval of signal positions between the start of the closest anchor at or before base *r*_*i*_ and the end of the closest anchor at or after base *r*_*i*_; this interval is extended by *bandwidth* positions on both ends (Figure 4).

**Figure 4:**
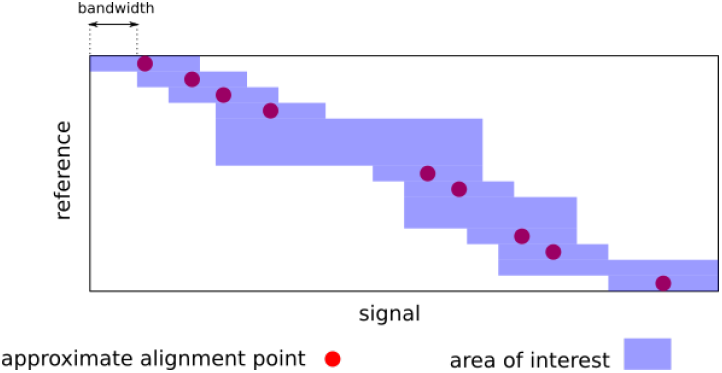
Areas of interest for efficient signal alignment

### 2.5. Signal Renormalization

Raw nanopore signal needs to be normalized by applying read-specific shift and scale parameters (denoted *α* and *β* respectively). Given a single raw signal level *s*_raw_ and the parameters *α* and *β*, the normalized signal level is (*s*_raw_*−α*)/*β*. This normalization can influence subsequent alignment [21] and other processing. In Nadavca, we initially normalize the signal by using the median signal value in a read as *α* and the median of |*s*_raw_*−α*| in a read as *β*. This median-based normalization is, however, sensitive to various nanopore data artefacts. For this reason, we improve the normalization by an iterative procedure [22, 21, 18].

In particular, after the initial normalization, the realignment algorithms assigns signal values to particular reference locations. For each such aligned location, our enhanced signal model predicts a particular expected signal level. In the next iteration, we update normalization parameters *α* and *β* to minimize the sum of squared errors between the expected and observed signal values in the read (omitting signal values aligned to transitional states). Subsequently, we repeat the process by re-computing the alignment and recomputing normalization again. We observed, that two iterations are sufficient to achieve a good accuracy.

### 2.6. Unsupervised Detection of DNA Modification Through Anomalies

DNA modifications typically cause small shifts in signal values. Given an accurate signal-to-reference alignment, this property can be exploited to detect DNA modifications by comparison of expected signal values predicted from a signal model to the observed signal values in individual events produced by the alignment. Here, we mostly follow the approach used by Tombo *de novo* modification detection module [18]. However, we substitute our improved signal-to-reference alignment and enhanced signal model to demonstrate their usefulness.

In particular, let [*a, b*] be an event aligned to the reference context *r*. Let *m* is the event mean, i.e. the mean of signal values *s*_*a*_ … *s*_*b*_. As an indicator of DNA modification, we compute the p-value that value *m* comes from a distribution *N*(μ(*r*), σ) defined by the enhanced signal model.

This simple method, however, does not lead immediately to accurate modification detection. On one hand, the shifts in signal values are small and thus individual p-values assigned to events are rarely significant. On the other hand, a typical modification affects several events near the modified base. Following the method outlined in Tombo, we combine p-values from three adjacent positions using the Fisher’s combined probability test [23] to form a more reliable indicator.

Finally, we output scores for positions where the reference sequence matches a user-supplied pattern recognized by the methyltransferase enzyme potentially active in a given sample. For each position with the pattern, we consider a window of length 11 centered on the potentially modified base and report the maximum score within this window.

### 2.7. Nanopore Sequencing

High-molecular weight (HMW) genomic DNA was prepared from an overnight culture of *Jaminaea angkorensis* C5b (CBS 10918; [24]) grown in yeast-peptone-dextrose (YPD) medium (1 % (w/v) yeast extract, 2 % (w/v) peptone, 2 % (w/v) glucose) at 28°C with constant aeration. Yeast cells were harvested by centrifugation, washed with 20 mM EDTA pH 8.0, resuspended in 100 mM EDTA pH 8.0, 2 % (v/v) 2-mercaptoethanol and incubated for 15 min at room temperature. Cells were then pelleted, resuspended in 0.15 M NaCl, 100 mM EDTA pH 8.0 and disrupted by vortexing with glass beads (0.4-0.5 mm, Sigma) three times for 30 s, followed by addition of sodium dodecyl sulfate (SDS) to 0.1 % (w/v). Nucleic acids were obtained by series of phenol and phenol : chloroform : isoamylalcohol (25 : 24 : 1) extractions, precipitated with an equal volume of 96 % (v/v) ethanol, washed with 70 % (v/v) ethanol, and air-dried. The precipitate was dissolved in TE buffer (10 mM Tris-HCl, 1 mM EDTA, pH 8.0) and RNA was removed by RNase A (150 *μ*g/mL digestion) for 30 min at 37°C. DNA was extracted by phenol : chloroform : isoamylalcohol (25 : 24 : 1), precipitated with 0.1 M NaCl and two volumes of 96 % (v/v) ethanol, washed with 70 % (v/v) ethanol, air-dried, dissolved in TE buffer and further purified using a Genomic-tip 100/G (Qiagen). HMW DNA of the yeast *Magnusiomyces capitatus* NRRL Y-17686 (CBS 197.35; [25]) was prepared essentially as described above, except that prior the phenol extractions, yeast cells were resuspended in 1 M Sorbitol, 10 mM EDTA pH 8.0 and converted to spheroplasts using zymolyase 20T (0.125 mg/mL; Seikagaku) treatment for 60-90 min at 37°C. Spheroplasts were then lyzed in 0.15 M NaCl, 100 mM EDTA pH 8.0, 0.1 % (w/v) SDS.

Nanopore sequencing was carried out in a MinION Mk1B device using a FLO-MIN106 (R9.4.1, revD) flow cell (Oxford Nanopore Technologies). The sequencing libraries were constructed essentially as described in the manufacturer’s instructions. Datasets of control unmodified DNA and DNA methylated *in vitro* were prepared using the PCR barcoding kit (SQK-PBK004, Oxford Nanopore Technologies). Briefly, HMW DNA was sheared in a g-Tube (Covaris) at 2700 ×*g* or 4300 ×*g* in a MiniSpin Plus centrifuge (Eppendorf). Fragmented DNA (*∼*200 ng) was treated using a NEBNext Ultra II End repair / dA-tailing module (New England Biolabs) and ligated to the Barcode adapters (BCA) from the sequencing kit (SQK-PBK004) using a Blunt/TA Ligase Master Mix (New England Biolabs). DNA samples (*∼*10 ng) were then amplified using the Rapid Barcode primers (LWB, SQK-PBK004) and LongAmp *Taq* DNA polymerase (New England Biolabs) and treated by exonuclease I (New England Biolabs). DNA aliquots (*∼*1 *μ*g) were modified using 160 *μ*M S-adenosyl methionine and methyltransferases M.*Eco*RI (40 U), M.*Bam*HI (12 U), or M.*Hha*I (25 U) (New England Biolabs) for 1 hour at 37°C. The methyl-transferase reactions were then stopped at 65°C for 20 min. The modification by M.*Taq*I (10 U) was carried out at 65°C for 1 hour. DNA was purified on AMPure XP beads and eluted into 10 mM Tris.Cl pH 8.0, 50 mM NaCl. Methylated and control barcoded DNAs were pooled in equal ratio. Rapid adapters (RAP, SQK-PBK004) were attached to the ends of pooled DNAs (*∼* 400 ng). The library preparation, flow cell priming, and loading were completed as described in the protocol (SQK-PBK004, version PBK_9073_v1_revA_23May2018).

### 2.8. Training and Testing Data Sets

All nanopore reads were base called by Albacore version

2.3.1. Base called reads were aligned to their respective reference genomes by BWA_MEM [26]. The reads that did not align to the reference for at least 80% of their length were discarded. Table 1 shows basic characteristics of resulting data sets.

**Table 1.**
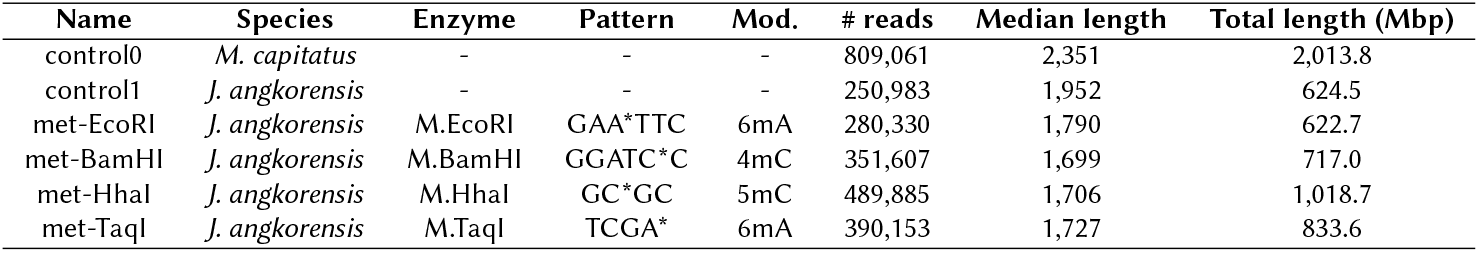
Data set overview. To produce data sets control0 and control1, the DNA was first synthesized using PCR and thus was free of any natural DNA modifications. For each of the met-* data sets, the DNA was further treated by one of the enzymes introducing specific DNA modifications at specific sites, as outlined in the table. The modification site within the pattern is marked with *.

Data set control0 was used for the purpose of training extended signal model. Mixtures of reads from control1 and met-* data sets were used for the purpose of testing the application of our methods to DNA methylation detection. Note that control0 and control1 data sets were produced from phylogenetically distant species and thus there is no overlap between training and testing sets.

## 3. Results

### 3.1. Realignment by Nadavca Eliminates Apparent Alignment Artefacts

Alignment of the signal to the reference sequence segments the signal into events, each event corresponding to a different context read by the pore. If the DNA moved through the pore at a constant speed, each event would span approximately 9 values.

We have used Tombo [18], Nanopolish [11], and Nadavca to realign the signal to the reference sequence, using data set control1. Figure 5 shows the comparison of event length distributions. Only Nanopolish allows events of length 0 (skip of the context), Tombo apparently has a minimum event length 3 (even though rarely it also outputs events of length 1 and 2), and in Nadavca we require each event to be of length at least 2. With Tombo, there is a clear bias against events of length 4, and preference for events of lengths 3 and 6; which appears to be an artefact of the alignment process. Interestingly, Nanopolish does not report events of length 1 or 2. In case of Nadavca, the only apparent artefact is a higher abundance of events of length 2, which is a consequence of the requirement of minimum length 2 for each event.

**Figure 5:**
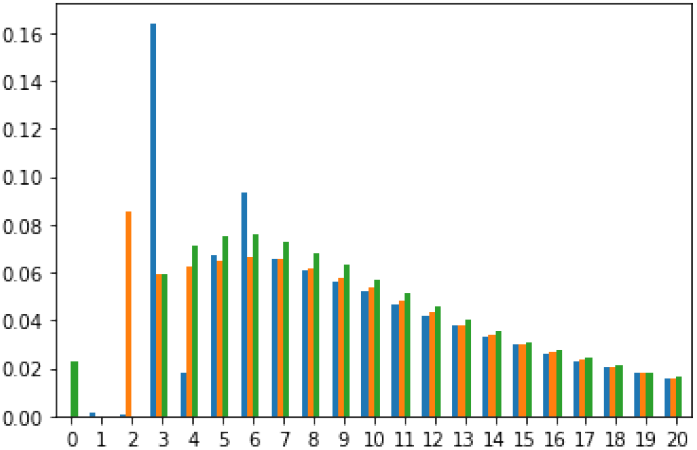
Event length distribution on dataset control1. Blue Tombo, orange Nadavca, green Nanopolish.

### 3.2. Unsupervised Single-Molecule Single-Site DNA Modification Detection

To evaluate the effect of Nadavca alignments on downstream analysis, we have implemented a simple tool nanometh for detection of methylation and other DNA modifications from the nanopore signals. We have used the following general framework for detection of modifications (see Methods for details):

1.Use Nadavca to align signal values of a read to the reference sequence

2.For each event implied by the alignment, compare the mean signal value within the event to the expected signal level from our enhanced signal model, producing a p-value

3.P-values from several adjacent positions are combined to a single score

4.Only scores for positions matching a user-supplied sequence pattern, such as GCGC, are written to the output.

We compare our results with de novo mode of Tombo. Note that unlike most other methods, Nadavca and Tombo do not require any training, besides training the sequence context model. To make the results comparable, we use similar formulas for computing the p-values and scores as Tombo, however, substituting different signal alignment and different signal context model. For Tombo we have used parameters supplied directly with the tool.

We use a newly produced testing set from the genome of fungus *Jaminaea angkorensis* [24]. Genomic DNA was amplified by PCR, producing DNA devoid of any native modifications, and then one part (control) was left unmodified, and each of four additional parts were modified by a different methyltransferase enzyme. Each enzyme methylates either adenine or cytosine at a fixed position within a specific sequence pattern (for example, M.HhaI methylates the first cytosine in the pattern GCGC). In each test, we use a mixture of reads from the control sample and from one of the modified samples. We consider only positions matching the pattern of the corresponding enzyme (in the reference genome) and classify them as positives or negatives based on the sample of origin. Note that the signal model was trained on a data set from a different organism to avoid overfitting.

We do not attempt to set a threshold for calling a site methylated, instead we compute AUC score which evaluates each method over all choices of the threshold. In the computation of accuracy, each pattern occurrence within each read is considered separately. Table 2 shows an overview of results and comparison to Tombo *de novo* mode. On all of our methylation data sets, nanometh significantly outperforms Tombo.

**Table 2.** Comparison of accuracy of methylation detection. The table lists AUC scores from the predictions of unsupervised tools (Tombo, Nadavca / nanometh) and a supervised tool trained for 5mC detection Nanopolish. Both Tombo and Nadavca report a score for each event, we aggregate these results at a particular position by using a maximum score in a window of 11 events centered around the site matching the methylation pattern. We switched off the horizontal aggregation of signal in Nanopolish (by setting minimum separator to 1), as it decreased its accuracy.

| Software tool | Parameters | Methylation pattern |  |  |  |
| --- | --- | --- | --- | --- | --- |
|  |  | GAA*TTC | GGATC*C | GC*GC | TCGA* |
| Tombo | *de novo* mode | 70.39 | 65.60 | 73.00 | 57.04 |
| Nadavca / nanometh |  | 75.03 | 68.09 | 78.59 | 63.03 |
| Nadavca / nanometh | offsets -1,0,2 | - | - | 90.8 | - |
| Nanopolish | min. separator 1 | - | - | 91.90 | - |

### 3.3. Comparison to Supervised Methylation Detection

Unlike Nadavca/nanometh, most tools for methylation detection are supervised, requiring a separate training set for each type of modification (usually obtained by creating artificially methylated samples). As a representative of supervised tools, we have selected Nanopolish, which also strives to find an accurate alignment of the signal values to the reference, but explicitly includes modified bases in an extended alphabet of its signal context model. We run Nanopolish with model parameters supplied with the tool.

Comparison of Nadavca to Nanopolish for the GCGC pattern is shown in table 2. There is still a large gap in performance, which is to be expected—while Nanopolish uses training data set to learn the differences between methylated and unmethylated bases, unsupervised tools only require information on composition of signal from unmodified bases.

We have also run Nadavca in a semi-supervised mode, where instead of aggregating information from a window of length 11, it is given an information on the most informative positions relative to the location of the pattern (in case of GCGC these are offsets -1, 0, and 2) and the final score is the sum of scores at these positions. In this mode, the performance of Nanopolish can be almost matched even with our otherwise unsupervised tool.

### 3.4. Contribution of Individual Algorithm Features

We were also interested, how individual features of Nadavca affect the accuracy. Therefore, we started with a baseline modification detection tool and added individual features one at a time: posterior alignment, renormalization, modeling transitional signal states, and using an extended k-mer model. The results are shown in Table 3. Replacing Tombo resquiggle with posterior alignment with renormalization already helps to increase the accuracy significantly. Adding transitions and extending the *k*-mer model further increases the accuracy, in some cases significantly.

**Table 3.** Contributions of individual features to methylation prediction accuracy.

| Squiggle alignment | Methylation pattern |  |  |  |
| --- | --- | --- | --- | --- |
|  | GAATTC | GGATCC | GCGC | TCGA |
| tombo (incl. renormalization) | 70.39 | 65.60 | 73.00 | 57.04 |
| nadavca (posterior alignment) | 66.23 | 65.19 | 71.60 | 58.60 |
| +renormalization | 73.90 | 64.65 | 80.65 | 61.01 |
| +transitions | 75.78 | 65.71 | 81.54 | 62.07 |
| +extended k-mer model | 75.03 | 68.09 | 78.59 | 63.03 |

## 4. Discussion

In this work, we introduced Nadavca, a nanopore signal aligner that incorporates several enhancements to the DTW algorithm. Compared to existing aligners, Nadavca’s output exhibits improved accuracy by eliminating length distribution artifacts and eliminating the need for event segmentation as a preliminary step. We demonstrated the efficacy of our alignment method by achieving state-of-the-art results in unsupervised methylation detection. Moreover, by incorporating additional information such as affected signal positions due to methylation, our approach yields comparable results to supervised tools.

In the future, we aim to further refine our alignment technique by addressing alignment artifacts, particularly in regions where the expected signals for two events are similar, resulting in ambiguous alignments.

## Funding

The research presented in this paper has been supported by funding from the Slovak Research and Development Agency grant APVV-18-0239 (JN), and by grants from Slovak Grant Agency VEGA 1/0234/23 (JN), 1/0463/20 (BB) and 1/0538/22 (TV). This research has also been supported by the Operational Program Integrated Infrastructure grant ITMS2014:313011ATL7.

## Acknowledgements

The strain of *J. angkorensis* was kindly provided by Matthias Sipiczki (University of Debrecen, Hungary). The strain of *M. capitatus* was kindly provided by Cletus P. Kurtzman and James Swezey (Agricultural Research Service, Peoria, IL, USA)

## Notes

### Competing Interest Statement

The authors have declared no competing interest.

